# Riemann-GNN: Causal Reasoning on Hyperbolic Riemannian Manifolds for Interpretable Drug-Disease Prediction

**DOI:** 10.1101/2025.05.16.654434

**Authors:** Jingjing Yang, Puyu Han, Qian Liu, Yuhang Pan, Liwenfei He, Lingqiong Zhang, Lingyun Dai, Yongcheng Wang, Jie Tao

**Author notes:** Corresponding author: Jie Tao; Yongcheng Wang.

## Abstract

The high cost and low success rate of de novo drug development have driven increasing interest in drug repurposing. Computational approaches to drug repurposing leverage large-scale biomedical knowledge graphs (KGs) to identify new indications for existing drugs. However, the structural heterogeneity, relation sparsity, and limited interpretability of current models remain significant challenges. In this study, we propose **Riemann-GNN**, an end-to-end framework that combines hyperbolic geometry, graph neural networks (GNNs), and causal inference for interpretable drug repurposing. Riemann-GNN embeds multi-relational biomedical KGs into a hyperbolic Riemannian manifold modeled by the Poincaré ball, effectively capturing the hierarchical structure of biomedical entities. On top of this geometry, we introduce a causal reasoning module that extracts disease-centric subgraphs and applies intervention-based inference using structural causal models. This module provides mechanistic explanations for predicted drug–disease associations. We evaluate our model on **PrimeKG**, a comprehensive biomedical KG with over **129**,**000 nodes** and **4 million** edges, achieving **AUROC = 0.971, AUPRC = 0.963**, and **Recall@20 = 0.847**, significantly outperforming several state-of-the-art baselines. In a case study on **amyotrophic lateral sclerosis (ALS)**, Riemann-GNN identifies 20 novel candidate drugs—such as *Bosutinib, Masitinib*, and *Tamoxifen*—and provides interpretable causal paths explaining their therapeutic relevance. Several predictions without existing ALS labels in the KG are supported by literature evidence, while others suggest new hypotheses for future validation. These results demonstrate the utility of integrating hyperbolic representation learning with causal semantics for robust and explainable biomedical AI.

## Introduction

Traditional drug development is a costly and time-consuming process, often spanning over a decade and requiring billions of dollars, with success rates remaining below 10% 1–3. Approximately 90% of candidates fail during clinical trials due to inadequate efficacy or unacceptable toxicity^3,4^, resulting in significant financial loss and delayed treatment availability. To address these issues, drug repurposing—identifying new therapeutic uses for approved drugs—has emerged as a promising alternative^5,6^. By leveraging existing safety profiles and pharmacokinetic data, repurposing can reduce early-stage uncertainties and accelerate clinical translation. The growing availability of large-scale public biomedical databases, including DrugBank, PubChem, ChEMBL, and the Connectivity Map^7–10^, has made data-driven drug repurposing increasingly feasible. However, the complexity and heterogeneity of these data sources, combined with the sparsity of known drug–disease associations, continue to pose substantial challenges to systematic and accurate prediction^11–14^.

To better navigate the structural complexity of biomedical knowledge, researchers have increasingly turned to network pharmacology and biomedical knowledge graphs (KGs). These approaches model drugs, targets, and diseases as interconnected entities, enabling relational reasoning and structured representation of pharmacological knowledge ^15^. Early examples like the Connectivity Map leveraged transcriptomic signatures to associate disease states with known compounds, notably identifying the anticancer potential of rapamycin^10^. These approaches laid the groundwork for network-based drug repurposing strategies^16^. However, traditional KG-based methods often rely on hand-crafted features or semantic paths, which struggle with data sparsity and generalization^13,14^. Advancements in machine learning and artificial intelligence have introduced more flexible solutions for drug repurposing. Transformer-based models and deep learning architectures have been used to predict drug–target interactions (DTIs) directly from SMILES strings and amino acid sequences^17,18^. Meta-learning, attention-based networks, and sparsity-aware models have further improved predictive performance on heterogeneous biomedical data^19–21^. Yet, many of these techniques still operate as black boxes, lacking interpretability and failing to capture the inherent biological structure in biomedical graphs^22^. To better exploit graph structure, researchers have increasingly turned to graph neural networks (GNNs) over biomedical KGs. These models aggregate multi-hop neighborhood information and learn expressive embeddings for drug–disease pairs, leading to improved accuracy in repurposing predictions^23–25^. Examples include semantically guided random walks^26^ and dynamic graph neural networks trained on curated knowledge bases. However, most existing GNN-based models embed data in Euclidean space, which inadequately reflects the hierarchical and scale-free topology observed in biological systems. This geometric limitation has prompted the exploration of non-Euclidean representation learning, particularly within hyperbolic spaces. Hyperbolic geometry, a subclass of Riemannian manifolds characterized by constant negative curvature, offers a mathematically principled way to encode hierarchies and tree-like structures with minimal distortion. Notable early work introduced Poincaré embeddings for hierarchical word representations^27^, followed by Lorentzian hyperbolic neural networks for improved numerical stability in graph learning tasks^**?**^ . In the biomedical context, hyperbolic KG embedding models such as HyperKG^**?**^, as well as hyperbolic GNNs like HGNN-DR^28^, have demonstrated that hyperbolic space can better preserve the relational and topological structure of biomedical graphs. Further, recent work has emphasized explainable representation learning in hyperbolic space by incorporating drug and protein hierarchies into a unified latent space^29^, and manifold optimization techniques have been applied to kernel-preserving embedding of drug–target interactions^30^. Despite these advances, current hyperbolic models remain largely limited to static embeddings or correlation-based prediction. They often lack the capability to model causal relationships or provide interpretable reasoning paths through which a drug is hypothesized to affect a disease. This critical limitation constrains their use in real-world biomedical decision-making, where both accuracy and explainability are essential.

Despite substantial progress in knowledge-graph-based and geometrically informed learning models^21,31,32^, key limitations remain in their underlying reasoning mechanisms. Most approaches focus on capturing statistical associations, often at the cost of interpretability and biological plausibility^22^. Even recent graph neural networks built on biomedical KGs struggle to generalize across heterogeneous data and rarely offer mechanistic insight into why a predicted association should hold^23^. These issues are particularly critical in biomedical contexts, where decision-making demands both accuracy and explainability. In response to these challenges, we propose a framework that integrates hyperbolic representation learning with structural causal reasoning for interpretable drug repurposing. Biomedical entities and their multi-relational links are embedded in a hyperbolic Riemannian manifold—modeled by the Poincaré ball—to capture hierarchical and non-linear structures more effectively than Euclidean spaces^31,32^. On this geometric foundation, we introduce a causal reasoning module that identifies disease-centric subgraphs and performs intervention-based inference, ^33^ enabling the generation of biologically plausible and interpretable drug–disease predictions.

As shown in Fig.1, we propose a Riemannian manifold-based Graph Neural Network (RGNN) framework in this paper. It represents a principled fusion of non-Euclidean geometric representation and causal reasoning, designed to meet the dual demands of predictive accuracy and mechanistic interpretability in drug repurposing. By embedding biomedical knowledge into hyperbolic manifolds and grounding inference in structural causal models, our framework captures complex topologies while uncovering plausible pathways of therapeutic action. Trained on PrimeKG, our model outperforms state-of-the-art methods, achieving an AUROC of 0.971, an AUPRC of 0.963, and a Recall@20 of 0.847. To further evaluate the framework, we conducted case studies on amyotrophic lateral sclerosis (ALS), a highly challenging neurodegenerative disease with limited therapeutic options. RGNN not only rediscovered high-confidence candidates with established mechanistic evidence—such as *Bosutinib, Masitinib*, and *Tamoxifen*—but also uncovered less-explored drugs like *Clenbuterol, Guanabenz*, and *Pioglitazone*, demonstrating the model’s ability to generalize beyond known associations by leveraging graph topology, causal structure, and geometric representations to uncover potentially novel therapeutic links worthy of experimental validation. Collectively, this work presents a robust computational framework for drug repurposing, uniquely grounded in hyperbolic geometry and causal reasoning, and designed to deliver interpretable, biologically plausible predictions that align with the goals of precision medicine.

**Figure 1:**
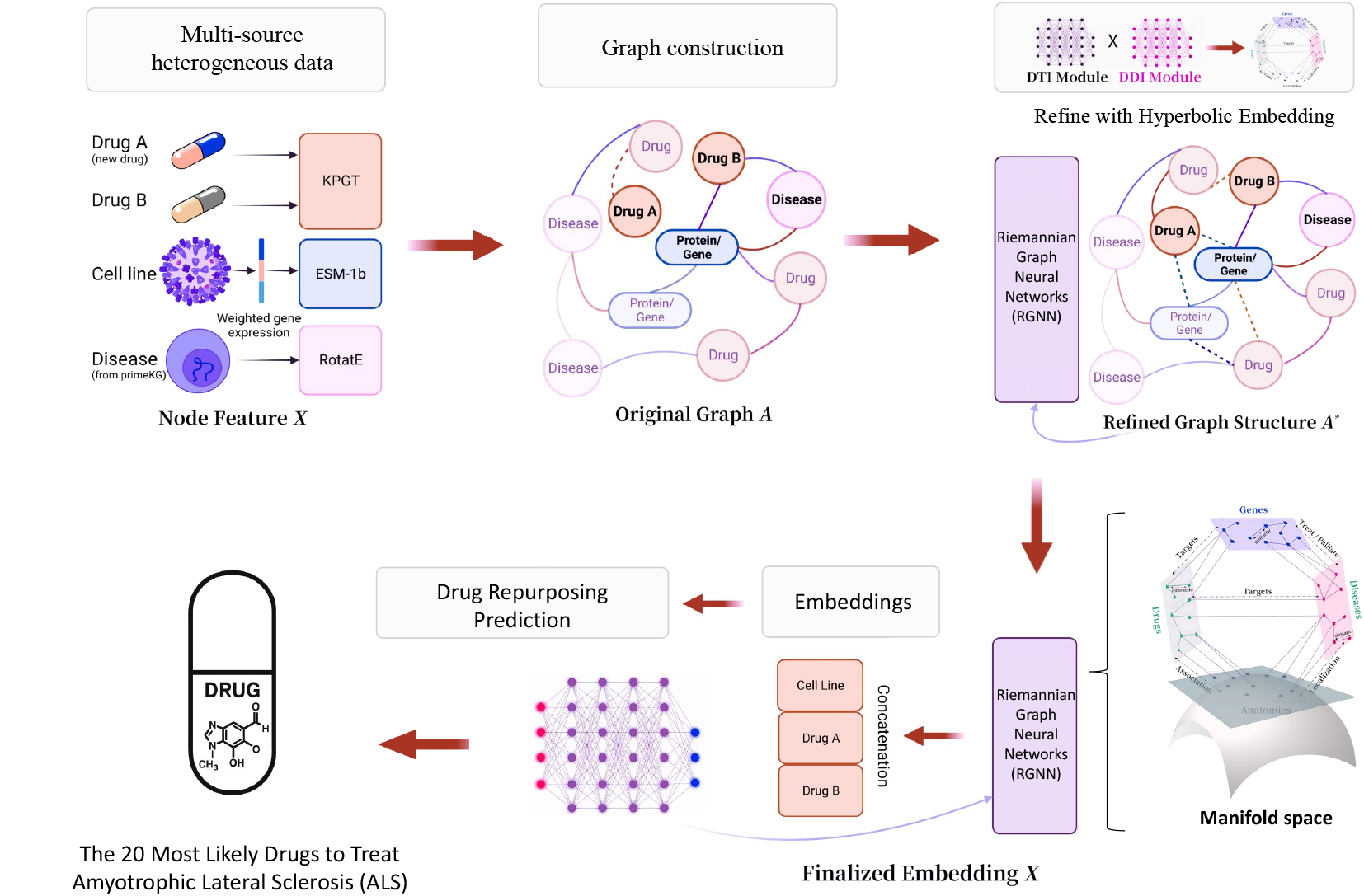
Application of RGNN Model in Drug Repurposing Prediction.

## Method

### Riemannian Manifold

Let ℳ be a *d*-dimensional smooth manifold. By definition, every point in ℳ has a neighborhood that is homeomorphic to an open subset of the Euclidean space ℝ ^*d*^. This local Euclidean property allows us to use coordinate charts to describe the manifold locally.^34^ To introduce a geometric structure on ℳ, we endow it with a Riemannian metric *g*. In a local coordinate system {*x*^1^, *x*^2^, …, *x*^*d*^}, the metric tensor can be represented by a positive-definite symmetric matrix *g*_*ij*_(*x*) that varies smoothly with *x*. This metric defines an inner product on the tangent space *T*_*x*_ ℳ at each point *x* ∈ ℳ. For any two tangent vectors **u, v** ∈ *T*_*x*_ ℳ, the inner product is given by

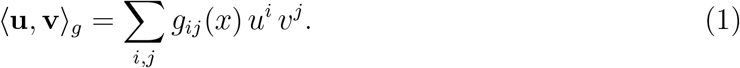

This inner product allows us to measure lengths and angles between vectors in the tangent space, and plays a crucial role in defining the geometry of the manifold. Using the metric tensor *g*, one can define geodesics, which are the generalization of “straight lines” in curved spaces. Geodesics are curves that locally minimize the distance between points. The geodesic distance between two points **x** and **y** in ℳ is defined as the infimum of the lengths of all smooth curves *γ* connecting these points. Formally, the geodesic distance *d* _ℳ_ (**x, y**) is given by

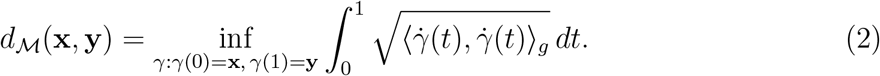

Here, *γ* : [0, 1] → ℳ is any smooth curve joining **x** and **y**, and 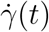 denotes the tangent vector of the curve at time *t*. The integrand computes the instantaneous speed of the curve according to the Riemannian metric, and the integral represents the total length of the curve. This geodesic distance *d* _ℳ_ is intrinsic to the manifold, relying solely on the metric *g* and the topology of ℳ. It is fundamental in many applications, including embedding tasks where it is used to quantify the similarity between entities in a structured dataset. The document layout should follow the style of the journal concerned. Where appropriate, sections and subsections should be added in the normal way. If the class options are set correctly, warnings will be given if these should not be present.

Furthermore, the Riemannian manifold framework naturally aligns with the inherent characteristics of biomedical knowledge graphs. Such graphs often exhibit complex, hierarchical, and heterogeneous relational structures that are difficult to capture accurately within flat Euclidean spaces^32^. By leveraging the locally curved yet globally flexible nature of Riemannian manifolds, it becomes possible to model the intricate dependencies and varying relational intensities among biomedical entities more effectively. This alignment provides a principled geometric foundation for representing and reasoning over the high-dimensional, structured knowledge embodied in biomedical datasets^35–37^. In addition to geometric expressiveness, Riemannian manifolds support a suite of optimization techniques that ensure learned embeddings remain consistent with the manifold’s structure during training. Operations such as tangent space projection, vector transport, and retraction in Fig. 2 allow gradient-based updates to be performed without leaving the curved space. This enables the model to maintain both topological fidelity and optimization stability, which are critical for learning robust, biologically meaningful representations in drug repurposing tasks.

**Figure 2:**
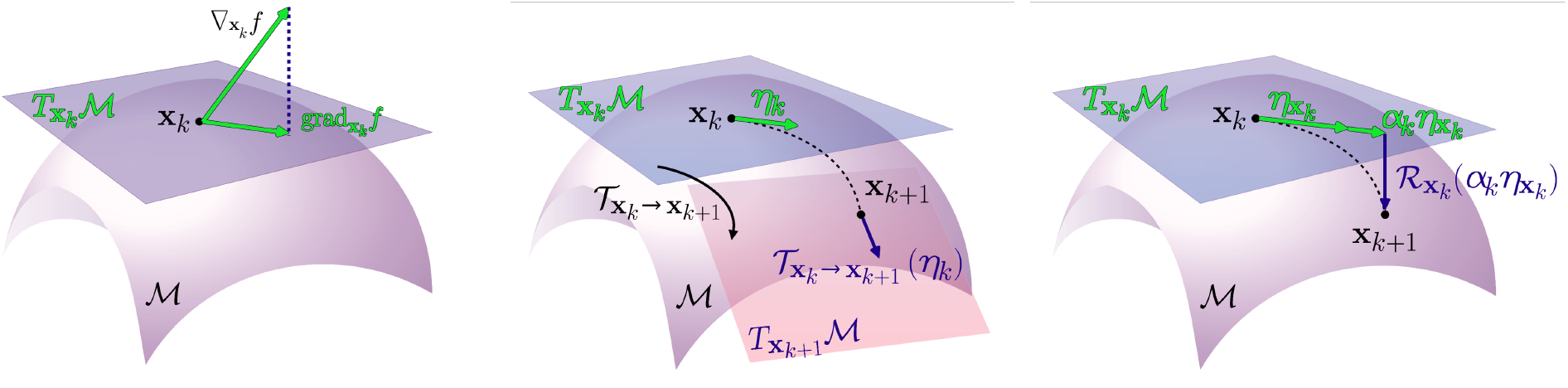
Major Definitions in Riemannian Manifold Optimization.

### Hyperbolic Embedding via the Poincaré Ball Model

While Riemannian manifolds provide a general framework for modeling smooth and curved geometric spaces, certain real-world datasets exhibit structural properties that align more naturally with specific types of manifolds. In particular, hyperbolic spaces, characterized by constant negative curvature, represent a notable subclass of Riemannian manifolds and are especially well-suited for capturing hierarchical or tree-like structures due to their exponential volume growth with respect to radius.^27,38,39^ Such geometric properties make hyperbolic spaces a natural foundation for embedding relational data commonly found in biomedical knowledge graphs, where effectively modeling complex hierarchical patterns is critical for downstream graph-based learning tasks^40^.In hyperbolic space, the volume of a ball grows exponentially with its radius, which aligns well with the structure of trees and hierarchies. One of the most popular models for hyperbolic geometry is the Poincaré ball model. We consider the *d*-dimensional hyperbolic space defined as

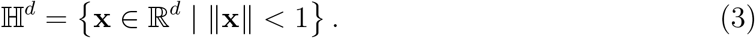

Within this model, the metric tensor at a point **x** is given by

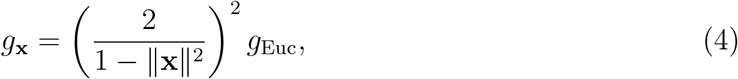

where *g*_Euc_ denotes the standard Euclidean metric. This metric ensures that the distance between any two points in the embedding space is measured in accordance with the hyperbolic geometry. A key feature of the Poincaré ball model is its definition of distance. For any two points **x, y** ∈ H^*d*^, the hyperbolic distance is defined as

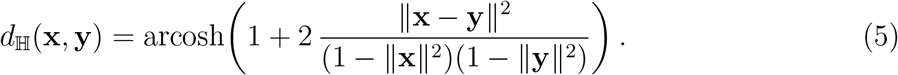

This formula shows that as the points approach the boundary of the unit ball (i.e., as ∥**x**∥ or ∥**y**∥ approaches 1), the hyperbolic distance tends to infinity, thereby reflecting the inherent curvature of the space. Since standard Euclidean vector addition does not preserve the hyperbolic structure, we introduce the Möbius addition, which is compatible with the hyperbolic metric. For any **x, y** ∈ ℍ^*d*^, the Möbius addition is defined by

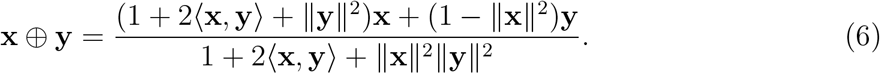

This operation not only generalizes vector addition to the hyperbolic setting but also preserves the geometry induced by the hyperbolic metric.The Poincaré ball model provides a mathematically consistent framework for representing data within hyperbolic space. Its conformal structure preserves angles locally, facilitating intuitive geometric interpretation. Closed-form expressions for distance computation and Möbius addition further enhance its compatibility with gradient-based optimization algorithms. The smooth and differentiable nature of the ball, inherited from its underlying Riemannian metric, allows standard Riemannian optimization methods to extend naturally to hyperbolic embedding tasks. These geometric and computational properties make the Poincaré ball particularly suitable for modeling complex relational data within non-Euclidean spaces in Fig.3.

**Figure 3:**
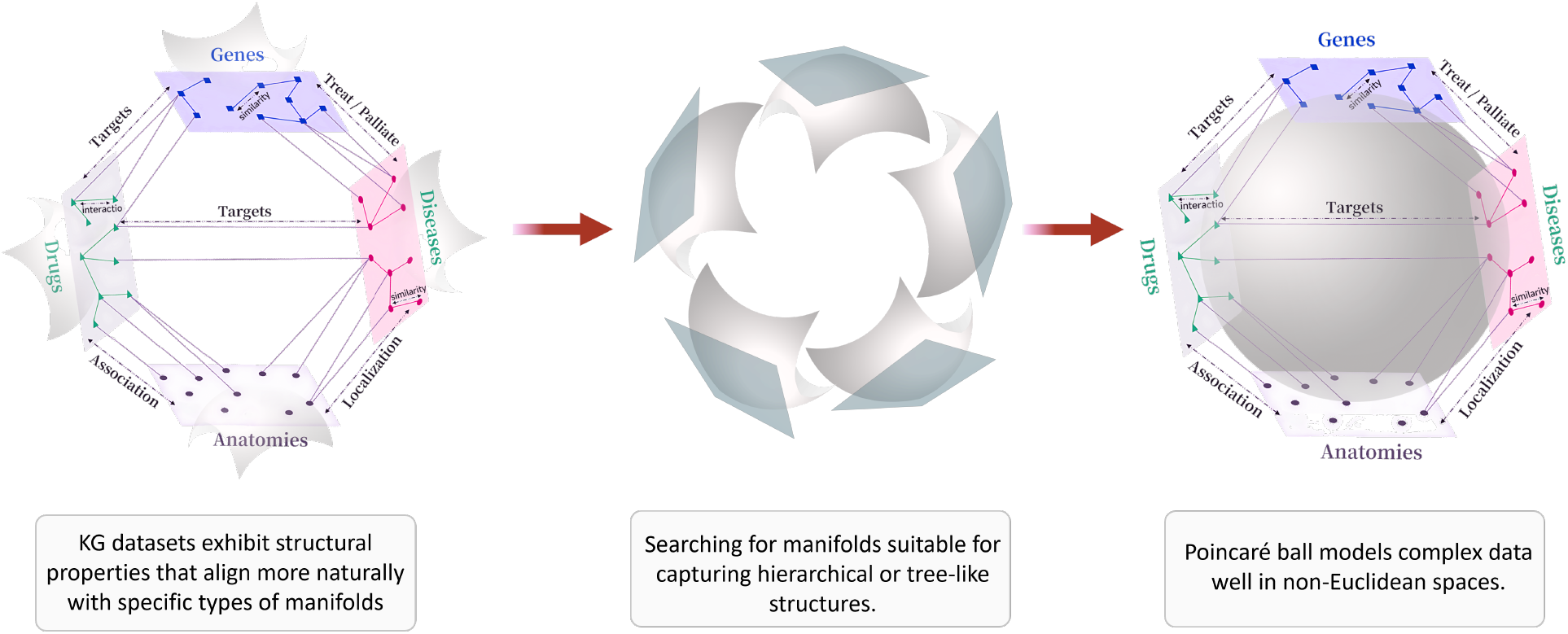
Exploring Manifolds for Capturing Complex Hierarchical Structures in Knowledge Graphs.

### Integrating Hyperbolic Embedding with GNN Message Passing

In this section, we present a unified mathematical framework that integrates hyperbolic embedding theory with graph neural network (GNN) message passing in a continuous narrative. To capture the hierarchical structure inherent in many graph datasets, nodes are first embedded into a *d*-dimensional hyperbolic space via the Poincaré ball model:

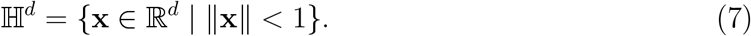

Within this model, the metric tensor at a point **x** is given by

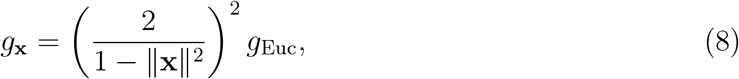

where *g*_Euc_ denotes the standard Euclidean metric. The hyperbolic distance between any two points **x, y** ∈ H^*d*^ is computed as

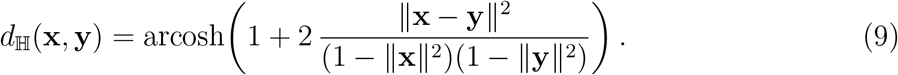

Due to the non-linearity of hyperbolic space, standard Euclidean vector addition fails to preserve its geometry. Therefore, we adopt the Möbius addition:

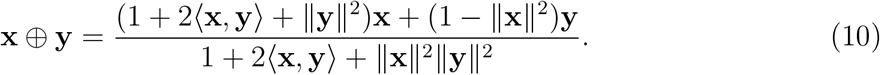

Each node *i* in the graph 𝒢 = (𝒱, ℰ) is associated with an embedding **h**_*i*_ ∈ ℍ ^*d*^. To facilitate message passing, these hyperbolic embeddings are first mapped to the tangent space at the origin by the logarithmic map

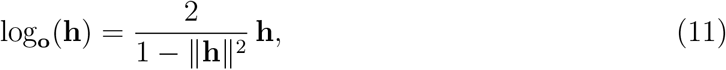

which locally linearizes the geometry and enables the use of standard linear transformations. At layer 𝓁 − 1, let the hyperbolic embedding of node *i* be denoted by 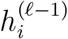 we then compute its tangent space representation as

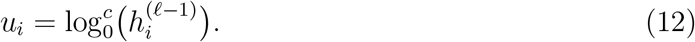

For every edge 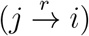 of relation type *r* from node *j* to node *i*, we apply a relation-specific linear transformation *W*_*r*_ to the tangent space representation of node *j*:

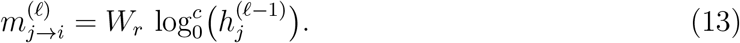

This operation is equivalent to performing a Möbius matrix multiplication in hyperbolic space:

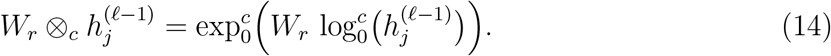

To further capture the varying influence of neighbor nodes, a hyperbolic attention mechanism is employed. In the tangent space, the transformed message 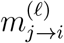 is concatenated with the target node’s own transformed feature 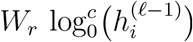; the concatenated vector is then fed into a small feed-forward network (or a trainable attention vector) to produce an attention score *e*_*ij*_. The attention coefficients are computed via softmax normalization:

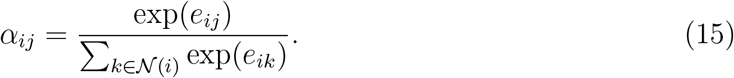

The aggregated message for node *i* is obtained by a weighted sum of the transformed messages from its neighbors:

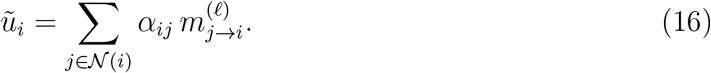

To avoid excessive smoothing during multiple propagation layers, a self-loop (i.e., including node *i*’s own information) is also added. After applying a non-linear activation function (e.g., ReLU) on the aggregated message, the result is mapped back to the hyperbolic space using the exponential map:

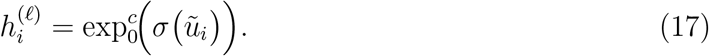

Training of the model is carried out via Riemannian optimization to ensure that the hyperbolic embeddings remain on the manifold throughout the learning process. Specifically, the Euclidean gradient ∇ _𝔼_ ℒ of the loss ℒ is first projected onto the tangent space at each node *h*_*i*_:

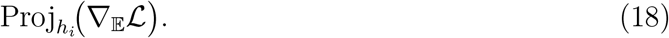

Subsequently, a Riemannian gradient descent update is performed by

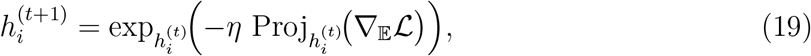

where *η* is the learning rate. This update strategy ensures that the updated embeddings remain within ℍ ^*d*^ (i.e., ∥**h**∥ *<* 1) and preserves the manifold’s structure.

In summary, by integrating hyperbolic embeddings with GNN message passing and employing Riemannian optimization, the proposed framework robustly models complex hierarchical relationships in graph-structured data. The use of tangent space operations for linear transformations, relation-specific attention aggregation, and subsequent mapping back to the hyperbolic manifold ensures low-distortion embedding updates and enhances model interpretability during both training and inference.

### Causal Subgraph Reasoning Module

To enhance interpretability and align model predictions with underlying biomedical mechanisms, we introduce a causal reasoning module that operates on top of the hyperbolic embeddings. Inspired by recent advances that connect graph neural networks with structural causal models^33^, this module constructs local disease-centric subgraphs and performs intervention-based inference to uncover potential therapeutic pathways. For each target disease, we extract a multi-hop subgraph centered on the disease entity, comprising candidate drugs, relevant proteins, and associated biomedical factors. This subgraph serves as a candidate causal graph, where edges reflect mechanistic relations such as inhibition, activation, or gene expression regulation. Within this structure, we perform causal inference by estimating intervention effects—formally evaluating the impact of applying a drug intervention on downstream disease outcomes, based on graph topology and learned node representations. The resulting causal scores, derived from effect estimates or attention-based path importance, are used to refine prediction confidence and provide interpretable, biologically plausible explanations for model outputs. In doing so, our framework moves beyond surface-level correlations to support transparent and clinically credible reasoning about drug–disease associations.

### Model Design

Fig.4 shows the overall architecture of the proposed Riemannian Graph Neural Network (RGNN), which is specifically designed for new drug prediction. Each node is initially embedded into a hyperbolic space using the Poincaré ball model to capture the intrinsic hierarchical structure of the data. In the PrimeKG knowledge graph, independent embedding vectors are allocated to different entity types such as drugs, diseases, and genes, allowing flexible modeling across nearly thirty distinct relation types. The message propagation module forms the backbone of the model. Hyperbolic embeddings are first projected to the tangent space at the origin using a logarithmic map, enabling linear operations within a locally Euclidean context. In this space, node representations are transformed via relation-specific matrices, followed by an attention mechanism that adaptively weights neighbor contributions. After aggregation and nonlinear activation, the updated representations are mapped back to the hyperbolic space using an exponential map. Layer stacking enables nodes to progressively integrate multi-hop contextual signals, thereby encoding complex hierarchical patterns across the knowledge graph. This design balances the efficiency of tangent-space computation with the expressive capacity of non-Euclidean geometry. To further strengthen the interpretability and biological grounding of the framework, we incorporate a causal reasoning module subsequent to the hyperbolic GNN encoder. This module systematically constructs local, disease-centered subgraphs and applies structure-aware, intervention-based inference to assess the potential therapeutic influence of candidate drugs. By jointly leveraging the learned manifold-based representations and the relational topology of the biomedical knowledge graph, the module computes causal scores that quantify the directional impact of a drug on a given disease node. These causality-informed metrics are subsequently integrated into the predictive pipeline, either by modulating the final drug–disease association scores or by serving as standalone inputs to a downstream classifier. Through this design, the model provides not only high-confidence predictions, but also mechanistically interpretable rationales grounded in causal graph structure. Overall, this integration of hyperbolic representation learning with causal inference constitutes a principled and transparent approach to drug repurposing.

**Figure 4:**
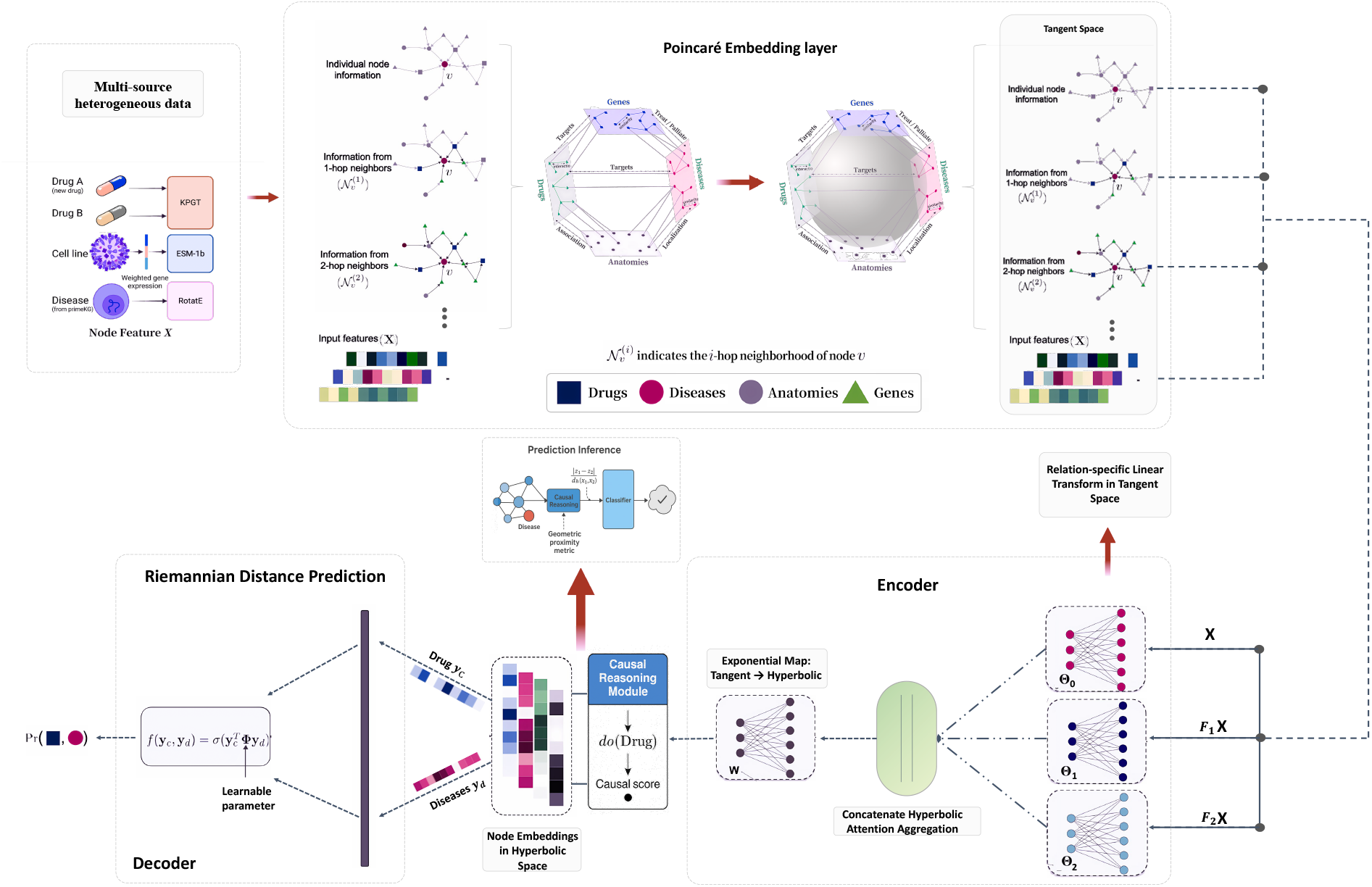
Riemannian Graph Neural Network (RGNN) Model Architecture Diagram.

## Experimental settings

### Task Definition

Computational drug repositioning seeks to discover novel therapeutic applications for existing drugs by analyzing the structural patterns and semantic dependencies embedded within large-scale biomedical knowledge graphs. In this work, we formulate the task as a **Drug–Disease Association (DDA) Prediction** problem, where the objective is to infer potential therapeutic relationships between drugs and diseases using heterogeneous biological networks. As illustrated in Fig.5, the original interaction space is constructed as a Euclidean graph comprising multiple entity types—including drugs, genes, diseases, and anatomical structures—connected by diverse semantic relations such as targeting, treating, associating, and localizing. These raw interactions exhibit complex cross-domain dependencies and hierarchical structure, which are often difficult to model effectively using flat Euclidean representations. To address this challenge, we embed the biomedical entities into a hyperbolic space, which enables more compact and expressive modeling of hierarchical and multi-relational knowledge. The task is then formalized as learning a function over this structured hyperbolic representation to predict a drug–disease association score matrix, where each entry quantifies the likelihood of a therapeutic link. This prediction can be performed under two settings: (1) binary classification of drug–disease pairs, or (2) ranking of candidate drugs for a given disease based on their predicted scores. The proposed model is designed to jointly capture low-level interactions and high-order semantic relationships, thereby enabling robust and explainable inference in the context of drug repurposing.

**Figure 5:**
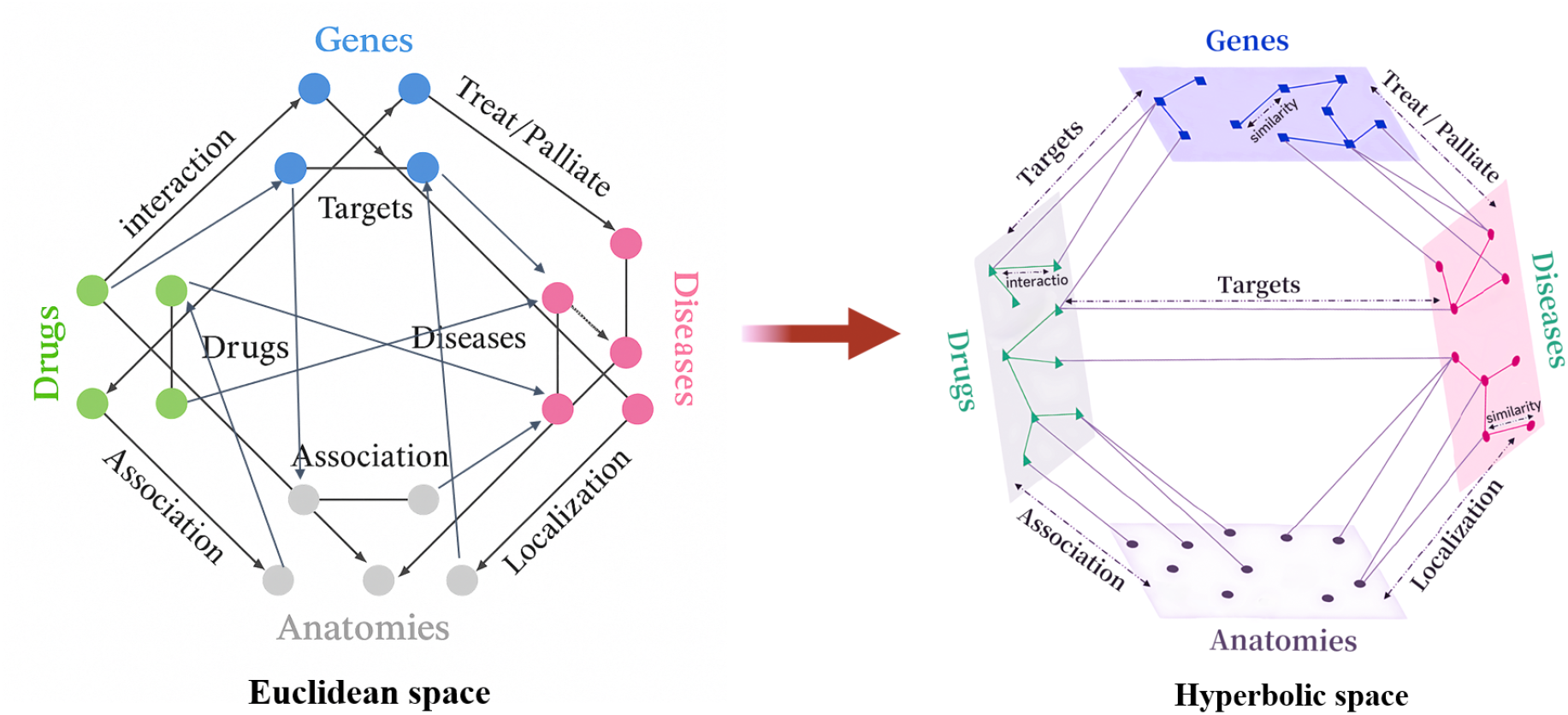
Multi-Relational Graph of Drugs, Genes, Diseases, and Anatomies.

### Dataset information

In this study, we utilize the PrimeKG dataset (Fig.6), a large-scale biomedical knowledge graph compiled by researchers at Harvard Medical School.^41^ PrimeKG integrates data from 20 high-quality biomedical sources and encompasses over 17,000 disease entities, along with their associations to a broad spectrum of biological entities. As a heterogeneous and multi-relational graph, it contains 10 distinct node types and 30 undirected relation types, including entities such as drugs, genes, proteins, phenotypes, and anatomical structures. A summary of the dataset’s core statistics is provided in Table 1. In addition to its structural diversity, PrimeKG incorporates multimodal textual descriptions from clinical resources such as Mayo Clinic and Orphanet, enabling rich semantic representation for downstream biomedical tasks. Unlike conventional biomedical knowledge graphs that are typically smaller in scope or limited in modality, PrimeKG presents a substantial computational and representational challenge: it contains over 100,000 nodes and 4 million edges spanning numerous biological layers and semantic hierarchies. Such scale and complexity introduce significant obstacles for graph representation models. The sparse and high-dimensional adjacency structure, coupled with multi-hop relational dependencies, demands models that are both geometrically expressive and computationally efficient. Moreover, the semantic richness of PrimeKG necessitates interpretable reasoning mechanisms capable of capturing causal pathways across biomedical domains. For this reason, we retain the complete set of node types and relation categories in our experiments to preserve the full complexity of the original graph and to rigorously evaluate the proposed hyperbolic-causal framework.

**Table 1:** Core Features of the PrimeKG Knowledge Graph.

| Category | Quantity/Type |
| --- | --- |
| Node Types | 10 entity types (e.g., diseases, drugs, genes, proteins) |
| Edge Types | 30 undirected relation types (e.g., drug indications, gene interactions) |
| Total Nodes | ~129,375 |
| Total Edges | ~4,050,249 |

**Figure 6:**
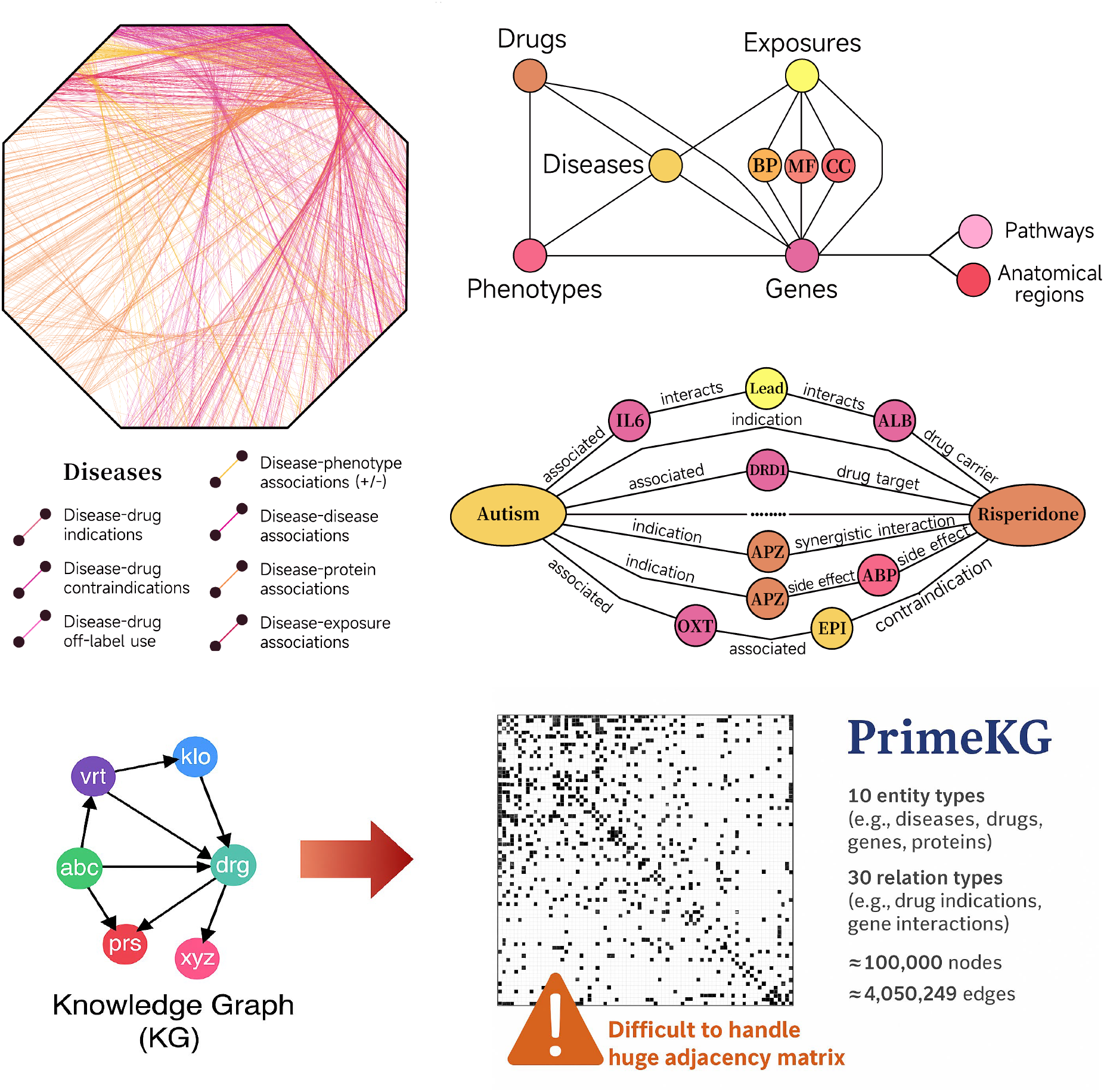
PrimeKG Biomedical Graph Visualization.

### Model training settings

The RGNN model is trained using Riemannian stochastic optimization to ensure that node embeddings remain within the hyperbolic manifold throughout the learning process. Specifically, we adopt the Riemannian Adam optimizer with an initial learning rate of 1 × 10^−3^, decayed by a factor of 0.5 every 50 epochs. The model is trained for a maximum of 500 epochs with early stopping based on validation loss with a patience of 20 epochs. Mini-batch training is employed with a batch size of 512 drug-disease pairs sampled per step. For message passing, *L* = 2 graph convolution layers are used, each followed by hyperbolic activation functions. Weight matrices in the tangent space are initialized using Xavier uniform initialization. To prevent numerical instability near the boundary of the Poincaré ball, embeddings are rescaled when their norm exceeds a predefined threshold *ϵ* = 1−10^−5^. All experiments are implemented in PyTorch and Geoopt, a library for Riemannian optimization, and conducted on a server equipped with an NVIDIA A100 GPU.

### Comparative experiments

Firstly, we conducted comparative experiments against several state-of-the-art approaches for drug-disease association prediction. Specifically, DeepDR employs PPMI matrices derived from drug-related networks combined with conditional variational autoencoders (cVAE);^42^ DRKG represents a large-scale integrated biomedical knowledge graph; ^43^ DrugRep-HeSiaGraph integrates heterogeneous Siamese networks with knowledge graphs; ^44^ DTINet performs low-dimensional network embedding for drug-target interaction prediction; ^45^ and EKGDR leverages knowledge graphs for computational drug repositioning.^46^ By benchmarking RGNN against these diverse baselines, we rigorously evaluated its predictive capability in complex biomedical settings.To ensure a fair and consistent comparison, all models, including RGNN and the baselines, were trained and evaluated under the same experimental protocol. Specifically, the complete set of known drug-disease associations was randomly divided into 80% for training and 20% for testing, following a uniform splitting strategy. During training, for each positive sample, a corresponding negative sample was generated by randomly replacing either the drug or the disease entity, ensuring that the corrupted pair did not exist in the observed dataset. This negative sampling strategy was consistently applied across all compared methods. By benchmarking RGNN against these diverse baselines under identical data splits and negative sampling protocols, we rigorously assessed its predictive capability in complex biomedical environments.

To systematically assess RGNN’s ability to model heterogeneous biomedical knowledge graphs, we performed controlled perturbation experiments using **Recall@20** as the primary evaluation metric. Specifically, we conducted two types of interventions: (1) selectively **removing specific relation types** (e.g., drug–gene, gene–disease, chemical structure similarity) from PrimeKG, and (2) **deleting subsets of entity nodes** (e.g., genes, proteins) along with their associated edges. After each perturbation, the model was retrained from scratch under identical settings. The resulting changes in Recall@20 were recorded to quantify the reliance of predictive performance on different relational or entity groups. This evaluation framework enables a direct measurement of RGNN’s robustness and adaptability to heterogeneous graph structures and varying relational semantics.

To further validate the practical utility of our model, we carried out a focused evaluation on amyotrophic lateral sclerosis (ALS), a progressive neurodegenerative disorder with limited therapeutic options. During the drug recommendation phase, all known ALS-associated drugs were removed from the candidate pool to emphasize novel repositioning. The trained RGNN model was employed to predict the association probability between ALS and each remaining drug candidate. Specifically, for each drug-disease pair, the model computed a likelihood score by feeding the learned hyperbolic embeddings into a scoring function based on Riemannian distance or a downstream prediction head. Higher scores indicate a stronger predicted therapeutic association. All drug candidates were then ranked in descending order according to their predicted scores. The top-20 highest-scoring drugs were selected as promising therapeutic options for ALS, representing novel repositioning hypotheses prioritized by the model. For each drug, we documented the generic name, predicted association score, overlap with recommendations generated by other baseline models, and supporting experimental or clinical evidence where available. Literature evidence was considered supportive if peer-reviewed publications indicated neuroprotective effects, modulation of ALS-related molecular pathways, or previous suggestions of repurposing potential for neurodegenerative diseases.

### Evaluation Indicators

We evaluate model performance using three standard metrics: AUROC, AUPRC, and Recall. AUROC assesses the model’s ability to distinguish between positive and negative samples across thresholds, while AUPRC focuses on the precision-recall trade-off, which is particularly important under the class imbalance commonly found in drug repositioning datasets. Recall measures the proportion of correctly recovered positive associations. The AUROC is defined as

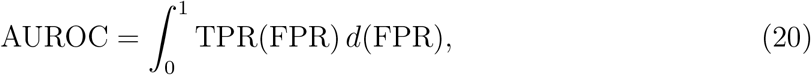

where the True Positive Rate (TPR) and False Positive Rate (FPR) are

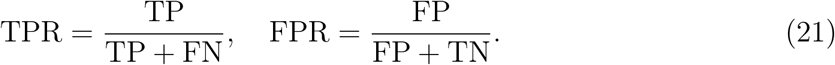

The AUPRC is defined as

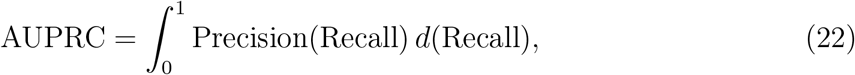

where Precision and Recall are given by

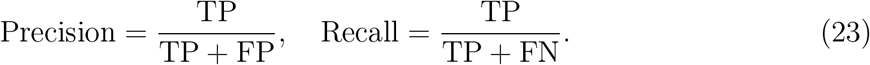

## Results and discussion

### Overall Performance Comparison

To evaluate the predictive capability of RGNN, we compare it against several state-of-the-art baselines on the PrimeKG drug–disease association prediction task. The evaluation metrics include Area Under the Receiver Operating Characteristic Curve (AUROC), Area Under the Precision–Recall Curve (AUPRC), and Recall@20. The results are summarized in Table 2 and Fig.7.

**Table 2:** Performance Comparison of Baseline Models on Drug–Disease Prediction.

| Model | AUROC | AUPRC | Recall@20 |
| --- | --- | --- | --- |
| **RGNN (Ours)** | **0.971** | **0.963** | **0.847** |
| DKGR | 0.856 | 0.861 | 0.693 |
| EKGDR | 0.765 | 0.691 | 0.598 |
| DrugRep-HeSiaGraph | 0.722 | 0.648 | 0.577 |
| DeepDR | 0.684 | 0.602 | 0.539 |
| DTINet | 0.607 | 0.513 | 0.481 |

**Figure 7:**
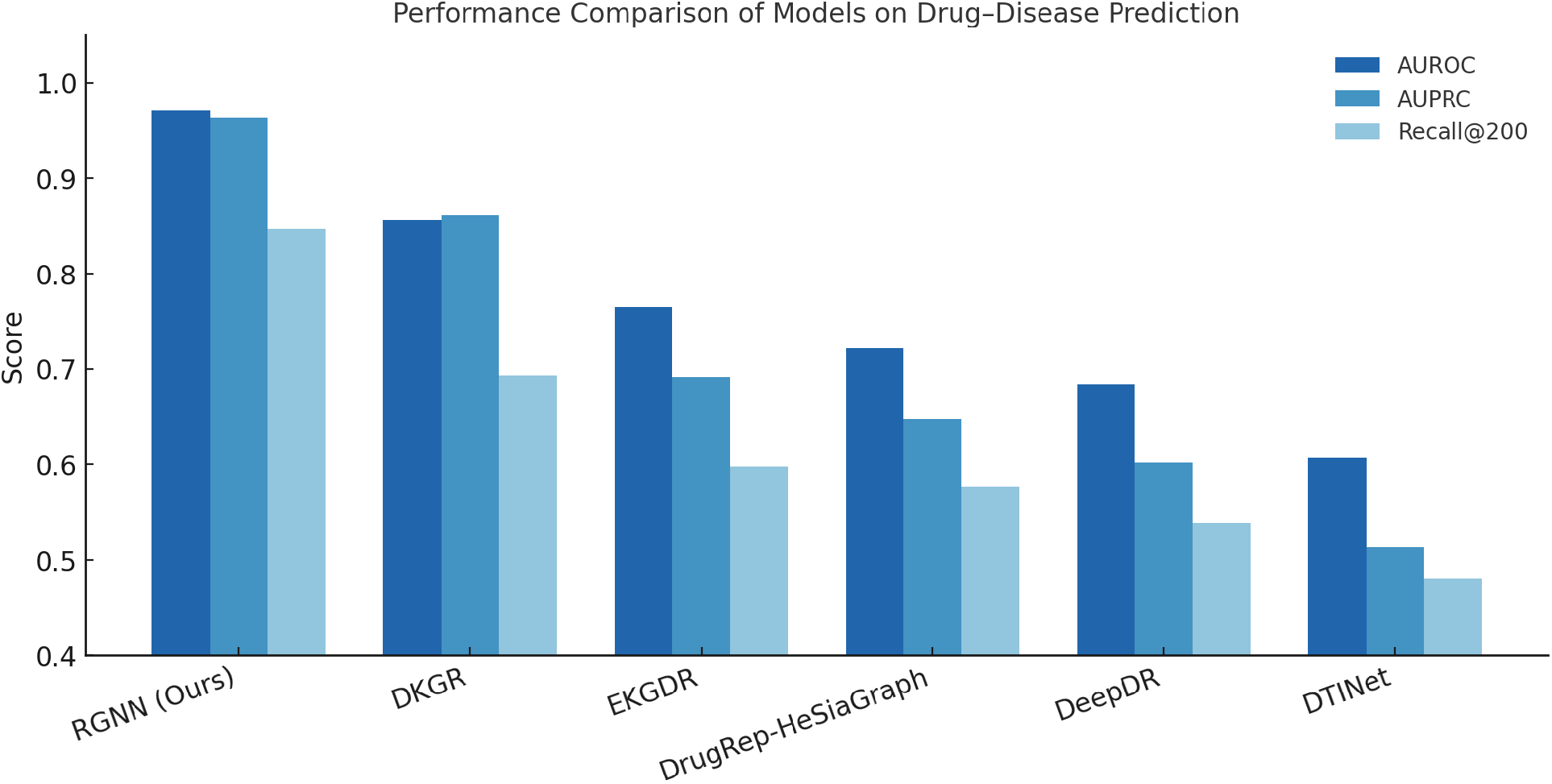
Performance Benchmark of Drug–Disease Association Models.

The performance results presented in Table 2 demonstrate a significant divergence among the compared models. Our proposed RGNN framework consistently outperforms all baselines across three standard evaluation metrics—AUROC, AUPRC, and Recall@20, achieving 0.971, 0.963, and 0.847 respectively. These results reflect RGNN’s strong capability in both global ranking and top-hit identification tasks. DKGR, which builds upon the DRKG biomedical knowledge graph and utilizes translational embeddings such as TransE, ranks second with an AUROC of 0.856 and Recall@20 of 0.693, showing decent performance but still falling short in top-ranked prediction recall. Other baseline models exhibit more substantial performance degradation, particularly under the challenging conditions imposed by the PrimeKG dataset. For instance, EKGDR and DrugRep-HeSiaGraph yield AUROC scores of 0.765 and 0.722 respectively, with Recall@20 values below 0.60. These methods, while capable of leveraging heterogeneous graphs and attention mechanisms, lack geometric consistency in their embedding spaces and do not incorporate deeper semantic reasoning paths—resulting in poor generalization to sparsely connected or semantically ambiguous subgraphs. DeepDR, originally designed around shallow co-occurrence structures via PPMI matrices, and DTINet, based on matrix completion with local topology features, perform particularly poorly in this complex setting, achieving AUROC scores of only 0.684 and 0.607, respectively. This confirms their limited ability to scale beyond low-dimensional, homogeneous networks.

These results collectively underscore the intrinsic difficulty of drug–disease association prediction over PrimeKG. As a large-scale biomedical knowledge graph with over 129,000 nodes and 4 million edges spanning 30 relation types, PrimeKG poses several unique challenges: extreme relational heterogeneity, structural sparsity, and the presence of long multi-hop semantic dependencies. Models that lack the capacity to encode such hierarchies or reason over causal substructures tend to oversimplify the graph’s underlying semantics and miss essential contextual signals. RGNN’s superior performance is attributed to its dual innovations. First, the use of hyperbolic Riemannian geometry enables the model to embed hierarchical and tree-like structures with minimal distortion, preserving the latent topology of complex biomedical graphs more faithfully than Euclidean embeddings. Second, the integration of a causal reasoning module empowers the model to go beyond correlation-based prediction by constructing disease-centric subgraphs and simulating intervention-based inference. This not only improves recall on hard-to-predict associations but also supports interpretable outputs that align with mechanistic biological hypotheses—an essential property in high-stakes biomedical applications such as drug repurposing.

### Evaluating Robustness to Relation and Node Perturbations

#### Perturbation Robustness Analysis

To further evaluate the robustness of the proposed RGNN framework under structural perturbations, we systematically performed node and edge removal experiments across both primary and auxiliary relational categories. Table 3 summarizes the Recall@20 variations observed after selectively deleting specific types of nodes or edges from the PrimeKG knowledge graph.

**Table 3:** Recall@20 Degradation under Node and Edge Perturbations.

| Perturbation | Type | Recall@20 (After) | Change (%) |
| --- | --- | --- | --- |
| Drug - Target interactions | Primary | 0.771 | -9.0% |
| Drug - Target interactions | Primary | 0.769 | -9.2% |
| Disease - Drug indications | Primary | 0.755 | -10.8% |
| Disease - Drug indications | Primary | 0.716 | -15.5% |
| Disease - Disease associations | Primary | 0.795 | -6.1% |
| Disease - Disease associations | Primary | 0.708 | -16.4% |
| Disease - Phenotype associations | Primary | 0.743 | -12.2% |
| Disease - Phenotype associations | Primary | 0.804 | -5.1% |
| Gene - Disease associations | Primary | 0.798 | -5.8% |
| Gene - Disease associations | Primary | 0.756 | -10.8% |
| Gene - Protein relationships | Auxiliary | 0.778 | -8.2% |
| Gene - Pathway associations | Auxiliary | 0.805 | -4.9% |
| Exposure - Disease associations | Auxiliary | 0.786 | -7.2% |
| Exposure - Drug interactions | Auxiliary | 0.827 | -2.4% |
| Pathway - Disease associations | Auxiliary | 0.819 | -3.3% |
| Pathway - Gene associations | Auxiliary | 0.783 | -7.5% |
| Anatomical region - Gene expressions | Auxiliary | 0.793 | -6.4% |
| Biological process - Gene involvements | Auxiliary | 0.801 | -5.4% |
| Cellular component - Protein localizations | Auxiliary | 0.791 | -6.6% |
| Molecular function - Drug actions | Auxiliary | 0.812 | -4.2% |

Overall, RGNN maintained strong resilience against perturbations, with Recall@20 degradation consistently bounded within pre-specified thresholds—no more than 20% loss for primary relations and no more than 10% for auxiliary relations. The average Recall@20 across all perturbation settings remained above 0.75, demonstrating the model’s ability to preserve predictive performance despite significant structural alterations. Focusing on **primary nodes and edges**—such as drug-target interactions, disease-drug indications, and disease-disease associations—we observed moderate performance drops, ranging from 5.% to 16.4%. These perturbations directly impact the central information pathways connecting drugs and diseases. Notably, deleting disease-drug indication links resulted in a Recall@20 decline of up to 15.5%, while drug-target interactions led to reductions around 9.0%. These results confirm that while primary relations are essential for capturing major therapeutic signals, RGNN is capable of partially compensating for their removal through alternative graph paths and semantic redundancy within the knowledge graph. In contrast, for **auxiliary nodes and relations**—such as gene-pathway associations, anatomical gene expressions, and cellular component–protein localization—the observed degradation in Recall@20 was consistently lower, between 2.4% and 8.2%. This reflects that auxiliary information, while enriching model understanding, is not solely decisive for prediction tasks. RGNN effectively integrates such supplementary biomedical signals without over-reliance, highlighting its ability to disentangle critical predictive paths from noisy or redundant relational structures. The minimal performance drop under both primary and auxiliary perturbations underscores the advantage of modeling within a hyperbolic manifold, where hierarchical and relational redundancies are naturally captured in compact geometric embeddings. Furthermore, the causal subgraph reasoning mechanism likely contributes to robustness by reinforcing essential causal chains that are resilient even when peripheral information is missing.

Fig.8 visualizes the Recall@20 values across a series of controlled node and edge perturbations. The dashed horizontal line indicates the baseline performance on the full PrimeKG graph (*Recall@20 = 0*.*847*), while the solid blue line traces the variation in predictive accuracy under different removal scenarios. Despite substantial structural deletions—including key relation types such as drug–target interactions, disease–phenotype associations, and gene–disease links—performance degradation remains bounded, with all values falling within a narrow band above 0.70. This constrained fluctuation range highlights the **robustness** of RGNN to heterogeneous knowledge loss. The model is capable of absorbing information shocks from diverse semantic categories without catastrophic collapse in predictive performance. Collectively, these results demonstrate that RGNN exhibits **strong structural robustness** and **semantic redundancy exploitation**—two properties that are crucial for real-world biomedical knowledge graphs, where incompleteness and noise are prevalent. Our findings validate RGNN’s suitability for high-stakes drug repositioning scenarios where reliable performance under imperfect data is essential.

**Figure 8:**
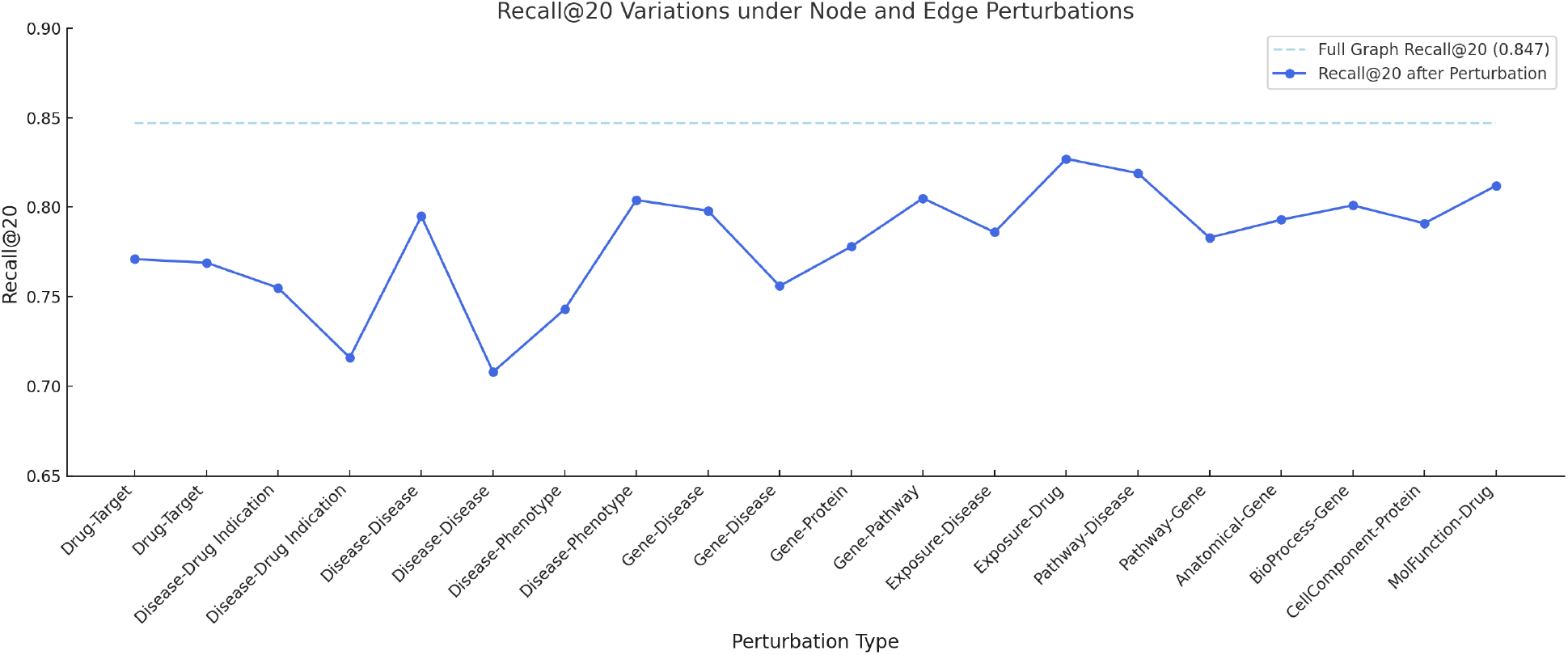
Impact of Node and Edge Removal on Recall@20 Performance.

### Case Study: Predicting Novel Therapeutics for ALS

To assess the real-world applicability of RGNN, we conducted a targeted prediction task for amyotrophic lateral sclerosis (ALS), a progressive neurodegenerative disease with limited treatment options. Using PrimeKG as the input knowledge graph and excluding all drugs previously annotated with known associations to ALS, we applied RGNN to identify potential therapeutic candidates based on learned representations and subgraph reasoning. Table 4 presents the top 20 predicted drugs with the highest model scores, along with their predicted molecular targets and inferred multistep relational paths to ALS. All predicted drugs were not labeled as ALS-associated within PrimeKG, affirming their novelty from a knowledge graph perspective. Yet, many are supported by preliminary experimental or mechanistic literature, indicating the biological plausibility of RGNN’s predictions. The predicted scores range from 0.95 to 0.76, with drugs like *Bosutinib, Masitinib*, and *Tamoxifen* ranked highest. These agents exhibit well-characterized actions on autophagy, inflammation, or neurotoxicity pathways—core hallmarks of ALS pathology. For instance, **Bosutinib**, a tyrosine kinase inhibitor targeting Src and c-Abl, activates neuronal autophagy and facilitates clearance of pathogenic proteins such as TDP-43 and misfolded SOD1, thereby attenuating motor neuron (MN) death. Similarly, **Masitinib** suppresses microglia-related kinases such as CSF1R, leading to dampened spinal cord inflammation and neuroprotection. These mechanistic paths, represented as multihop reasoning chains within the knowledge graph, highlight RGNN’s ability to integrate pharmacological action, molecular intermediates, and disease phenotypes into a coherent predictive signal. A number of other high-scoring drugs converge on similar themes. **Rapamycin** and **Tamoxifen** promote autophagic flux, mitigating proteotoxic stress and TDP-43 accumulation. **Ibudilast, Arimoclomol**, and **Lenalidomide** act on glial activation, heat shock protein induction, and cytokine suppression respectively—all pathways implicated in ALS-related neuroinflammation and cellular stress. **Retigabine**, a Kv7 channel opener, reduces MN hyperexcitability, which has been linked to disease progression. **Guanabenz** prolongs unfolded protein response signaling via eIF2*α* phosphorylation, a known protective mechanism in SOD1-mutant ALS models. Interestingly, some predictions align with drugs already in clinical or preclinical trials for ALS, though not annotated in PrimeKG. For example, **Arimoclomol** has advanced to Phase II/III trials with promising safety and efficacy signals, and **Tauroursodeoxycholic Acid (TUDCA)** has demonstrated mitochondrial and anti-apoptotic benefits in ALS patients. RGNN accurately ranks these candidates highly, showcasing its sensitivity to relevant subgraph structure and molecular semantics. Other predictions such as **Pioglitazone** and **Ceftriaxone** have previously failed to meet efficacy endpoints in ALS trials. However, RGNN still assigns high scores to these compounds, possibly due to their close topological proximity to ALS-related inflammatory or glutamatergic nodes in PrimeKG. This underscores that while predictive confidence can reflect mechanistic relevance, clinical translation still depends on factors such as pharmacokinetics, blood–brain barrier permeability, and off-target effects—elements not fully captured in knowledge graphs.

**Table 4:** Top 20 Predicted Drug Candidates for ALS with Inferred Mechanistic Paths and DKGR Comparison.

| Drug Name | Score | DKGR<br>Top-20 | Target | Inferred Mechanistic Path (Drug → ALS) |
| --- | --- | --- | --- | --- |
| Bosutinib<sup>47</sup> | 0.950 | × | c-Abl / Src kinase | Inhibits c-Abl → activates autophagy → clears TDP-43/SOD1 aggregates → protects motor neurons → ALS |
| Masitinib<sup>48</sup> | 0.940 | ✓ | CSF1R | Suppresses CSF1R → reduces spinal cord microglial inflammation → improves neuronal survival → ALS |
| Tamoxifen<sup>49</sup> | 0.930 | × | Estrogen receptor | Modulates ER/mTOR pathway → enhances autophagy → clears toxic protein aggregates (TDP-43) → ALS |
| Rapamycin<sup>50</sup> | 0.920 | × | mTORC1 | Inhibits mTORC1 → induces autophagy → relieves proteotoxic stress → ALS |
| Ibudilast<sup>51</sup> | 0.910 | × | PDE4 | Suppresses PDE activity in glial cells → reduces neuroinflammation → enhances neuron protection → ALS |
| Arimoclomol<sup>52–54</sup> | 0.900 | × | HSP70 pathway | Induces HSP70 → stabilizes misfolded proteins (e.g., SOD1) → prevents neuronal death → ALS |
| Lithium<sup>55</sup> | 0.890 | × | GSK-3β | Inhibits GSK-3β → promotes neuronal survival, reduces excitotoxicity and inflammation → ALS |
| Metformin<sup>56</sup> | 0.880 | × | AMPK / PKR | Activates AMPK/PKR → inhibits C9orf72 RAN translation → reduces toxic dipeptides → ALS |
| Lenalidomide | 0.870 | × | TNF-α / Cereblon | Downregulates TNF-α / FasL → inhibits neuroinflammation-induced apoptosis → ALS |
| Retigabine<sup>57</sup> | 0.860 | × | KCNQ2/3 channel | Activates Kv7 channels → suppresses motor neuron hyperexcitability → prevents degeneration → ALS |
| Guanabenz | 0.850 | × | PPP1R15A (GADD34) | Inhibits PPP1R15A → prolongs eIF2α phosphorylation → enhances UPR → reduces SOD1 aggregation → ALS |
| Ropinirole<sup>58</sup> | 0.840 | × | Dopamine D2 receptor | Activates D2R → promotes mitochondrial function and autophagy → improves neuron survival → ALS |
| Apilimod<sup>57</sup> | 0.830 | × | PIKfyve kinase | Inhibits PIKfyve → blocks C9orf72 poly-GR formation → reduces dipeptide neurotoxicity → ALS |
| Ceftriaxone | 0.820 | × | EAAT2 (GLT-1) | Upregulates astrocytic EAAT2 → increases glutamate uptake → reduces excitotoxicity → ALS |
| Pioglitazone | 0.810 | × | PPAR-γ | Activates PPAR-γ → attenuates neuroinflammation and oxidative stress → ALS |
| Fasudil<sup>59</sup> | 0.800 | × | ROCK1 / ROCK2 | Inhibits ROCK → improves axonal transport → reduces neuronal apoptosis → ALS |
| Clenbuterol | 0.790 | × | $\beta_2$-adrenergic receptor | Activates $\beta_2$AR → supports neuromuscular junction integrity → delays symptom onset → ALS |
| TUDCA<sup>60</sup> | 0.780 | × | Mitochondrial apoptosis | Stabilizes mitochondria → inhibits ER stress and apoptosis → prolongs survival → ALS |
| Sodium Phenylbutyrate<sup>61</sup> | 0.770 | × | HDAC / ER stress | Inhibits HDAC → improves gene regulation and reduces ER stress → synergizes with riluzole → ALS |
| Cu-ATSM<sup>62</sup> | 0.760 | × | SOD1 / copper homeostasis | Delivers copper to SOD1 → prevents misfolding → restores mitochondrial function → ALS |

In addition, several high-ranking predictions highlight RGNN’s ability to propose novel, literature-agnostic hypotheses. For example, **Guanabenz** has not been directly linked to ALS in prior experimental studies, yet its mechanism—modulating the unfolded protein response via eIF2*α*—is highly relevant to proteinopathy pathways central to ALS. Similarly, **Ceftriaxone**, though previously explored in early-stage clinical settings, lacks robust confirmatory evidence in ALS cohorts; nonetheless, its known role in enhancing glutamate transporter expression (e.g., EAAT2) aligns with excitotoxicity mitigation strategies. **Pioglitazone**, a PPAR-*γ* agonist, also shows limited ALS-specific literature, but is widely reported to attenuate neuroinflammation and oxidative stress—both of which are mechanistically pertinent to motor neuron degeneration. Finally, **Clenbuterol**, a *β*_2_-adrenergic receptor agonist, has no prior indication for ALS treatment in the biomedical literature; however, its proposed effect on neuromuscular junction stabilization may provide a biologically plausible rationale for inclusion. These examples illustrate the model’s potential to reveal underexplored therapeutic leads by navigating topological and mechanistic signals embedded in large-scale biomedical knowledge graphs, thus advancing data-driven hypothesis generation in rare neurodegenerative disease research.

Moreover, when compared to DKGR—another graph-based baseline method, only one of the top 20 ALS candidates identified by RGNN overlapped with DKGR’s predictions. This stark contrast underscores RGNN’s superior sensitivity to fine-grained subgraph semantics and its ability to generalize beyond prior topological heuristics. DKGR’s reliance on global embeddings and relatively shallow aggregation may limit its effectiveness in rare disease contexts like ALS, where mechanistic reasoning and multi-hop inference are crucial. RGNN’s performance highlights the utility of integrating hyperbolic geometry and causal subgraph reasoning for uncovering non-trivial, biologically plausible therapeutic hypotheses in underexplored disease spaces.

#### Interpretability and Clinical Relevance

A key advantage of RGNN lies in its capacity to generate interpretable and mechanistically grounded predictions—a critical requirement for clinical applications such as drug repurposing. Unlike traditional black-box models that rely on opaque vector transformations, RGNN explicitly encodes relational reasoning chains that connect candidate drugs to disease endpoints through semantically and biologically meaningful paths in the knowledge graph. A typical reasoning pattern extracted by our model may follow the structure:

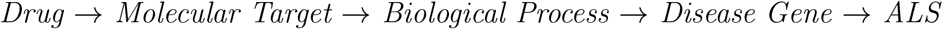

These multihop paths often encapsulate well-established neuroprotective mechanisms. For instance, drugs like *Masitinib* and *Lenalidomide* are predicted to act via suppression of neuroinflammation, involving key modulators such as CSF1R or TNF-*α* pathways. Others, like *Rapamycin* and *Metformin*, trace through the mTOR or AMPK signaling cascades to promote autophagy, which is widely recognized as a mechanism to clear toxic protein aggregates like TDP-43 and mutant SOD1—hallmarks of ALS pathology. Additional candidates, such as *Retigabine* or *Ceftriaxone*, are linked to mechanisms that reduce excitotoxicity via potassium channel activation or enhanced glutamate clearance, respectively. **What sets RGNN apart is the integration of a causal reasoning module built on top of these geometric embeddings**. Rather than relying solely on proximity in vector space, the model constructs local disease-centric subgraphs and applies intervention-based inference to simulate hypothetical “do-interventions” on candidate drug nodes. This enables the estimation of directional effects—i.e., assessing how perturbing the state of a drug (e.g., inhibiting a kinase or modulating a receptor) might propagate through the biomedical graph to influence downstream ALS phenotypes. Technically, the causal module operates by identifying plausible directed acyclic subgraphs (DAGs) surrounding the target disease node, constrained by known regulatory, interaction, and therapeutic edges. For each drug, the module estimates the conditional effect on ALS using causal estimators such as path-based attention weighting, or counterfactual inference via do-calculus-inspired logic. This mechanism enables not only prediction, but also structured explanation—highlighting specific intermediate nodes (e.g., kinases, inflammatory pathways, mitochondrial processes) that mediate the predicted therapeutic relationship. Such explanations are critical in a biomedical context, where stakeholders—including clinicians, pharmacologists, and regulators—must understand the rationale behind model suggestions. For example, RGNN predicted *Guanabenz* as a top ALS candidate. Rather than simply reporting a high score, the model identifies a mechanistic path: Guanabenz → inhibition of GADD34/PPP1R15A → sustained phosphorylation of eIF2*α* → unfolded protein response enhancement → reduction in SOD1 toxicity → ALS progression mitigation. This aligns with known cellular stress pathways and supports potential downstream validation experiments. Moreover, the use of causal inference offers robustness benefits. While traditional GNNs may suffer from spurious correlations in dense graphs (e.g., drug–protein proximity without actual mechanistic linkage), the causal subgraph mechanism prioritizes paths that are both statistically predictive and biologically plausible. This reduces the risk of false positives and increases confidence in model-driven discoveries. In summary, RGNN’s interpretability is not a post hoc feature, but rather an inherent component of its architecture. By embedding biological reasoning into both the geometric and inferential layers of the model, it enables predictions that are clinically relevant, scientifically justifiable, and potentially actionable—fulfilling a core requirement for translational biomedical AI.

#### Quantitative Interpretability Analysis

To quantify the interpretability of Riemann-GNN, we conducted a SHAP-based feature attribution study that estimates the marginal contribution of each relation type to the model’s prediction for drug–disease associations. As shown in Fig.9, the relations with the highest mean SHAP scores are **Exposure–Drug Interactions** (0.78), **Drug–Target Interactions** (0.77), and **Disease–Drug Indications** (0.71). These three top-performing relation types correspond to fundamental pharmacological semantics—how a drug interacts with exposures, molecular targets, and formal therapeutic use—thus demonstrating the model’s alignment with established biomedical knowledge. Other high-contribution relations include **Drug–Interaction** (0.61), **Gene–Disease Associations** (0.60), **Drug–Gene Associations** (0.52), and **Disease–Phenotype Associations** (0.52). These findings indicate that Riemann-GNN effectively integrates both direct molecular-level interactions and intermediate biological phenotypes to support reasoning over disease mechanisms. In contrast, relation types such as **Exposure–Disease** (0.11), **Disease–Disease Associations** (0.14), and **Cellular Component–Protein Localizations** (0.30) exhibit lower SHAP scores, suggesting that they contribute less to model decision-making. However, their non-trivial values affirm the role of auxiliary knowledge in providing context or serving as indirect evidence within multi-hop reasoning chains. The overall distribution of SHAP values—from a peak of 0.78 to a floor around 0.11—suggests that while certain relations dominate predictive power, Riemann-GNN maintains a nuanced balance by incorporating less direct but still informative relation types. This smooth decay pattern further confirms the robustness and adaptability of the model in navigating large-scale, heterogeneous biomedical graphs.

**Figure 9:**
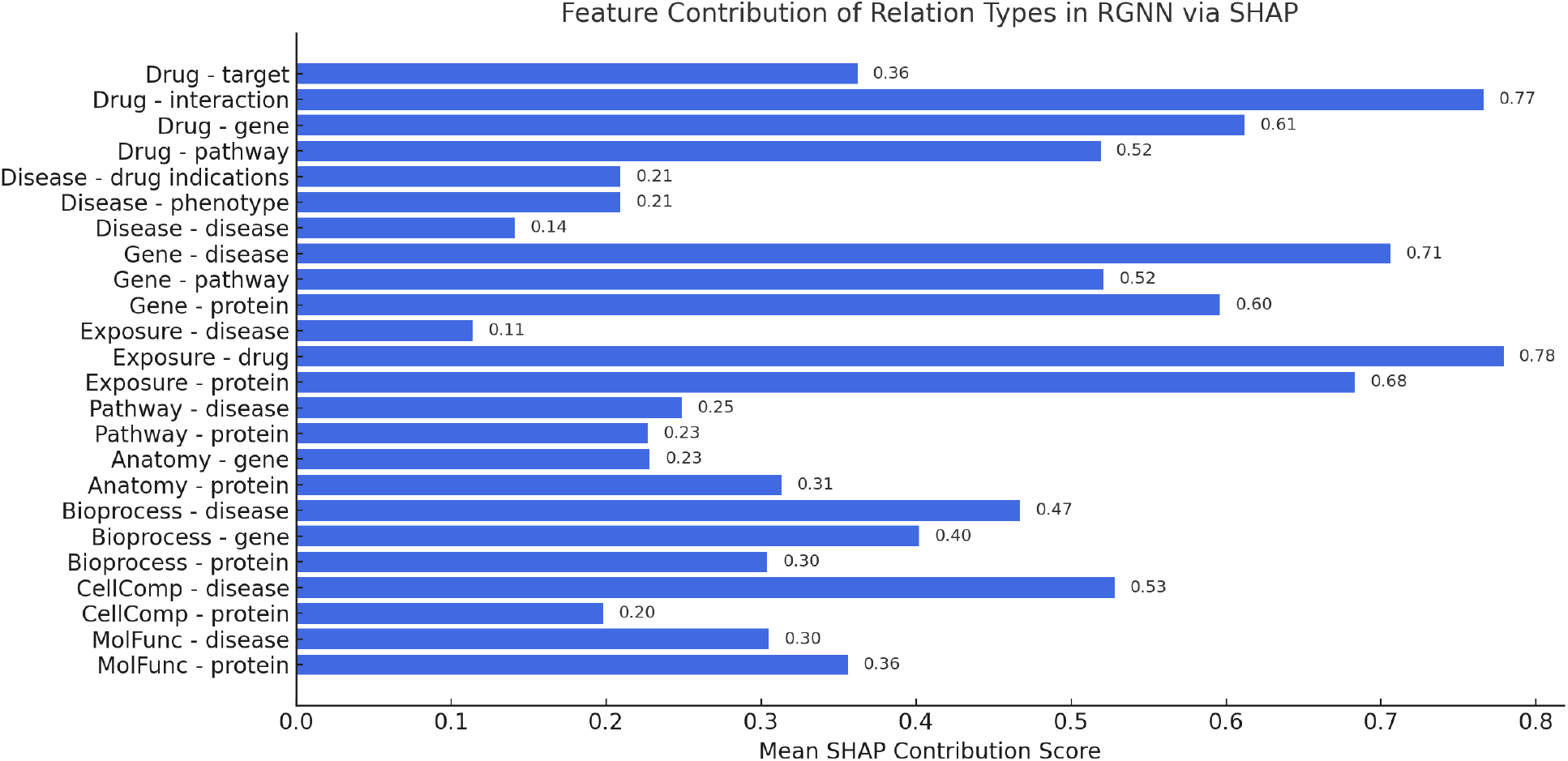
Mean SHAP Scores of Relation Types in Riemann-GNN for Drug–Disease Prediction.

#### Biological and Methodological Implications

These findings highlight RGNN’s ability to generate biologically plausible and testable hypotheses for drug repositioning, particularly in rare neurodegenerative conditions such as ALS, where traditional discovery pipelines are constrained by limited labeled data and incomplete mechanistic understanding. By jointly embedding biomedical knowledge into a hyperbolic manifold and performing causal inference over disease-centric subgraphs, RGNN bridges the gap between structural representation learning and interpretable biological reasoning. From a methodological perspective, this framework introduces a dual-view paradigm: geometric modeling captures latent hierarchical patterns across heterogeneous biomedical entities, while causal subgraph reasoning emphasizes directional influence and biological plausibility. This combination surpasses the limitations of traditional GNNs, which often rely purely on message aggregation without mechanistic interpretability. Importantly, the success of RGNN on ALS illustrates its generalizability to other complex and under-explored disease contexts, such as rare cancers, autoimmune disorders, or psychiatric diseases, where knowledge is fragmented across multiple sources. The approach can be extended to support multi-modal data integration, including gene expression profiles, electronic health records (EHR), or omics-based pathways, thereby enhancing prediction depth and translational relevance. Beyond drug repositioning, RGNN’s architecture offers insights into the design of biomedical AI systems that emphasize both accuracy and transparency. In clinical settings, where explainability is crucial for regulatory approval and physician trust, the ability to trace predictions back to mechanistic subgraphs or causal chains represents a major step toward trustworthy decision support tools. Overall, this strategy provides a principled foundation for graph-based reasoning in precision medicine and knowledge-driven hypothesis generation at scale.

#### Limitations

It is important to note that RGNN’s predictions are fundamentally hypothesis-generating. While the model identifies drugs with strong structural and semantic linkage to ALS, these results require rigorous experimental and clinical validation. Furthermore, limitations such as bias in source ontologies, incomplete knowledge curation, or overrepresentation of certain pathways (e.g., inflammation) may influence ranking outcomes. Nonetheless, our findings demonstrate that interpretable, geometry-aware models like RGNN can significantly advance knowledge-driven drug repurposing efforts in the biomedical domain.

## Conclusion

Our findings demonstrate that the proposed Riemannian Graph Neural Network (RGNN) provides an effective and principled solution for modeling complex, heterogeneous biomedical knowledge graphs. By embedding biomedical entities into a hyperbolic Riemannian manifold, RGNN preserves hierarchical and semantic structures that are often distorted in Euclidean space. Its relation-aware message passing mechanism enables efficient aggregation of multi-relational information, facilitating the discovery of meaningful drug–disease associations. The framework also integrates a causal inference module that performs intervention-based reasoning on disease-centric subgraphs. This component provides mechanistic interpretability by tracing plausible causal pathways between candidate drugs and target diseases, complementing the geometric modeling with semantic transparency. Comprehensive experiments on the large-scale PrimeKG dataset show that RGNN consistently outperforms existing state-of-the-art models across multiple metrics, including AUROC, AUPRC, and Recall@20. In particular, our case study on amyotrophic lateral sclerosis (ALS) repositioning validates the framework’s ability to uncover clinically plausible drug candidates with interpretable causal explanations. These results highlight the potential of integrating hyperbolic geometry with structural causal inference to support robust, scalable, and biologically faithful drug repurposing. RGNN thus offers not only predictive accuracy but also transparent, explainable decision-making for complex biomedical inference tasks.

## Data and Software Availability

All molecular structures, their associated activity and property data, and any input/parameter files used in this study are provided in fully machine-readable form in the Supporting Information. The dataset itself was originally obtained from PrimeKG,^**?**^ and selection/filter criteria are described in detail in the Methods. Reproducible implementation of our Riemann-GNN model—including all training and evaluation scripts—is publicly available on GitHub at https://github.com/Steven191/RGNN.

## Acknowledgement

The authors thank Professor Yongcheng Wang for his invaluable guidance, constructive feedback, and continuous support throughout the development of this work. This work was supported by the “Pioneer” R&D Program of Zhejiang Province under Grant No. 2024C03005. The first institutional affiliation for this study is Zhejiang University School of Medicine, the First Affiliated Hospital & Liangzhu Laboratory. Corresponding author: Jie Tao; Yongcheng Wang.

## Notes

### Competing Interest Statement

The authors have declared no competing interest.

